# Immunization by replication-competent controlled herpesvirus vectors

**DOI:** 10.1101/299230

**Authors:** David C. Bloom, Robert K. Tran, Joyce Feller, Richard Voellmy

## Abstract

Replication-competent controlled virus vectors were derived from virulent HSV-1 wildtype strain 17*syn*+ by placing one or two replication-essential genes under the stringent control of a gene switch that is co-activated by heat and an antiprogestin. Upon activation of the gene switch, the vectors replicate in infected cells with an efficacy that approaches that of the wildtype virus from which they were derived. Essentially no replication occurs in the absence of activation. When administered to mice, localized application of a transient heat treatment in the presence of systemic antiprogestin results in efficient but limited virus replication at the site of administration. The immunogenicity of these viral vectors was tested in a mouse footpad lethal challenge model. Unactivated viral vectors - which may be regarded as equivalents of inactivated vaccines - induced detectable protection against lethality caused by wildtype virus challenge. Single activation of the viral vectors at the site of administration (rear footpads) greatly enhanced protective immune responses, and second immunization resulted in complete protection. Once activated vectors also induced far better neutralizing antibody and HSV-1-specific T cells responses than unactivated vectors. To find out whether the immunogenicity of a heterologous antigen was also enhanced in the context of efficient transient vector replication, a virus vector constitutively expressing an equine influenza virus hemagglutinin was constructed. Immunization of mice with this recombinant induced detectable antibody-mediated neutralization of equine influenza virus as well as a hemagglutinin-specific T cell response. Single activation of viral replication resulted in a several-fold enhancement of this immune response.

**IMPORTANCE:** We hypothesized that vigorous replication of a pathogen may be critical for eliciting the most potent and balanced immune response against it. Hence, attenuation/inactivation (as in conventional vaccines) should be avoided. Instead, necessary safety should be provided by placing replication of the pathogen under stringent control and of activating time-limited replication of the pathogen strictly in an administration region in which pathology cannot develop. Immunization will then occur in the context of highly efficient pathogen replication and uncompromised safety. We found that localized activation in mice of efficient but limited replication of a replication-competent controlled herpesvirus vector resulted in a greatly enhanced immune response to the virus or an expressed heterologous antigen. This finding supports the above hypothesis as well as suggests that the vectors may be promising novel agents worth exploring for the prevention/mitigation of infectious diseases for which efficient vaccination is lacking, in particular in immunocompromised patients.

## INTRODUCTION

Vaccination has been spectacularly successful, but there remain important infections for which therapeutic or preventative vaccines either do not exist or are of insufficient efficacy. One such example are infections caused by the herpes simplex viruses HSV-1 and HSV-2. Recent estimates have the world-wide prevalence of HSV-1 at 67% and of HSV-2 at 11% (1,2). Perinatal herpes infection as well as infection in immune-compromised persons can be life-threatening. HSV infection has many complications, resulting in significant morbidity and even mortality. Currently, there is no vaccine for treating or preventing HSV-1 or HSV-2 infections. There are many ongoing research projects directed toward developing effective vaccines, of which some are being/have been tested in the clinic (3,4), with GEN-003 (Genocea) (5), HSV529 (Sanofi) (6) and HSV-2 0ΔNLS (Rational Vaccines) (7) receiving the most attention. Another example are influenza infections. Annual infection rates for influenza are between 5 and 20%, and the yearly death rate worldwide lies between 250,000 and 500,000 according to WHO estimates. At this time, the vaccines remain seasonal, i.e., they need to be updated/reformulated for every influenza season. Typical subunit vaccines are at least trivalent, comprising hemagglutinins from H1N1 and H3N2 influenza A strains and an influenza B strain (referred to as TIV). More recently, live attenuated influenza virus vaccines (LAIV) were introduced. An updated systematic review and meta-analysis published in 2012 documented a mean efficacy for TIV of 62% against RT-PCR or virally confirmed influenza (all age groups) (8). Limited data were available for adults 65 years of age and older. Mean efficacy for LAIV in young children was higher, but little or no protection was found in adults (18 years and older). A recent study reported a mean overall vaccine efficacy of 48% for the 2015-2016 season, and lower efficacies for persons 50 years of age and older (9). LAIV was ineffective in this season, even in children. Notable is a study on young children that was carried on over two seasons in both of which there was a good match between vaccine (TIV) and circulating strains (10). Vaccine efficacy was 66% in the first season and −7% in the second season. Thus, currently available influenza vaccines provide moderate protection against virologically confirmed disease, but protection is not long-lasting and is especially tenuous in the senior population. Ongoing preclinical and clinical work to develop better influenza vaccines includes formulation of adjuvanted vaccines, MVA-vectored T cell vaccines expressing conserved viral proteins, and hemagglutinin stalk-based vaccines for obtaining broadly protective antibody responses (11-13). A third example concerns *Mycobacterium tuberculosis* infections that affect about one third of the world population with yearly death rates of over one million. Of concern in the industrialized world is the rise of drug-resistant forms of the disease (14). The only vaccine in use is the (live) Bacillus Calmette-Guérin (BCG) vaccine developed in 1921 whose efficacy appears to be variable in adults. Candidate vaccines currently under development include virally vectored vaccines presenting BCG antigens, adjuvanted subunit vaccines and whole-cell vaccines based on *M. tuberculosis* or closely related mycobacterial species (15-17). Further examples include malaria and HIV/AIDS. As these examples reveal, adequate protection against many important infections is not yet available. Research continues, and new immunization approaches appear to be urgently needed.

The question whether live attenuated vaccines that have retained some capacity for replication provide more robust protective immunity than inactivated non-replicating vaccines was already debated more than 50 years ago. In the case of poliomyelitis vaccines, i.e., the live attenuated vaccine of Sabin vs. the inactivated vaccine of Salk, and also in the case of measles vaccines, a live attenuated vaccine based on the Edmonston B strain vs. an inactivated vaccine, the live attenuated vaccines eventually gained the upper hand (although safety concerns result in the predominant use of the Salk vaccine in regions in which the disease is considered eradicated) (18-21). It is noted that the latter historical debates were not conclusive because the vaccines compared differed in more than just their capacity to replicate (different routes of administration, modification of surface antigens by the inactivation process, etc.). Several more recent studies readdressed the issue in a more rigorous fashion. One such study compared immune responses to an HIV envelope antigen expressed from an attenuated replicating and a replication-defective adenovirus vector in a chimpanzee model (22). Inoculation with the replicating recombinant resulted in a greater frequency of HIV envelope-specific interferon-gamma-secreting peripheral blood lymphocytes, better priming of T cell proliferative responses, higher anti-envelope binding and neutralizing antibody titers and better antibody-dependent cellular cytotoxicity. In another study, an attenuated vaccinia virus (MVTT) expressing the spike protein (S) of SARS-CoV (MVTT-S) was compared with a replication-defective vaccinia virus (MVA) expressing the same protein (MVA-S). Intranasal or intraoral vaccination of mice with MVTT-S produced 20-100-fold higher neutralizing antibody levels than MVA-S (23,24). Yet another study employed a mouse ocular challenge model to demonstrate that an attenuated replicating HSV-1 strain (ICP0^−^) but not a replication-defective HSV-1 strain (ICP4^−^) elicited a protective immune response (25). These findings support the theory that attenuated replicating viruses induce more complete and more potent immune responses to autologous or heterologous antigens than corresponding non-replicating viruses.

We hypothesized that a virus vector that could replicate in a controlled fashion with nearly the same efficiency as the respective wildtype virus (referred to herein as “replication-competent controlled virus vector”) would induce an even more potent and complete immune response to itself or an expressed heterologous antigen than a corresponding attenuated vector (26). Our hypothesis is in part based on the rational expectation that an efficiently replicating virus will produce a stronger inflammatory response than an attenuated virus, which inflammatory response will result in a potent activation of the innate immune system and, consequently, in strong and lasting B and T cell responses (27). To realize such an immunization strategy, a regulation system must be employed that reliably and stringently controls viral replication as well as is capable of being turned on and off at will. However, virus disseminates after administration. Simply restricting the number of replication cycles will not be enough: depending on the number of cycles allowed, there will be a more or less pronounced manifestation of the toxicity typical for the viral vector used. For example, allowing an HSV-1 vector to replicate for a certain period of time may cause unacceptable neurotoxicity. Hence, the regulation system must be capable of exerting regional control over viral replication so that the immunizing virus only replicates in a locale in which it is certain not to cause a disease phenotype.

To facilitate construction of a replication-competent controlled virus vector, advantage may be taken of gene switches developed for eukaryotic cells (28). This choice implies that the backbone virus must be selected from the group of viruses having a double-stranded DNA genome (e.g., herpes viruses) or producing a double-stranded DNA intermediate during replication and utilizing the host machinery for transcription. It appears that the above-described requirements are best met by promoters of certain heat shock (HSP) genes that are stringently controlled by reversibly heat-activated endogenous transcription factor HSF1. A preferred choice may be the promoter of the human HSP70B (HSPA7) gene, which promoter has been found to be highly inducible and to exhibit very low basal expression wherever this has been tested (28-30). A virus in which one or, better, two replication-essential genes are placed under the control of this promoter could be activated to undergo one round of localized replication by administration of a mild heat dose to the inoculation region. Further rounds may be induced by repeat heat treatment. However, virus replication may be inefficient because heat activation of HSP promoters is rapidly reversible as well as may be unsafe because unintended weak activation could be triggered by high fever, intoxication, etc., in an inoculated subject. We previously described a two-component HSP promoter-based gene switch that lacks the latter unwanted properties (31) (Fig. 1A). The first component comprises a gene for a small-molecule regulator (SMR)-activated transactivator (TA). The TA gene is controlled by a promoter cassette containing an HSP70B promoter and a TA-responsive promoter (TRP). The second component is a TRP promoter for driving a selected viral gene (VG). Heat treatment prompts expression of the TA that is inactive unless activated by its SMR. Activated TA mediates expression of the viral gene as well as autoactivates its own expression. The gene switch is inactivated by withdrawal of the SMR. The present report presents a first study of immune responses induced by replication-competent controlled HSV-1 vectors in which one or two replication-essential genes are subjected to regulation by such a heat- and SMR-dependent gene switch.

**FIG 1.**
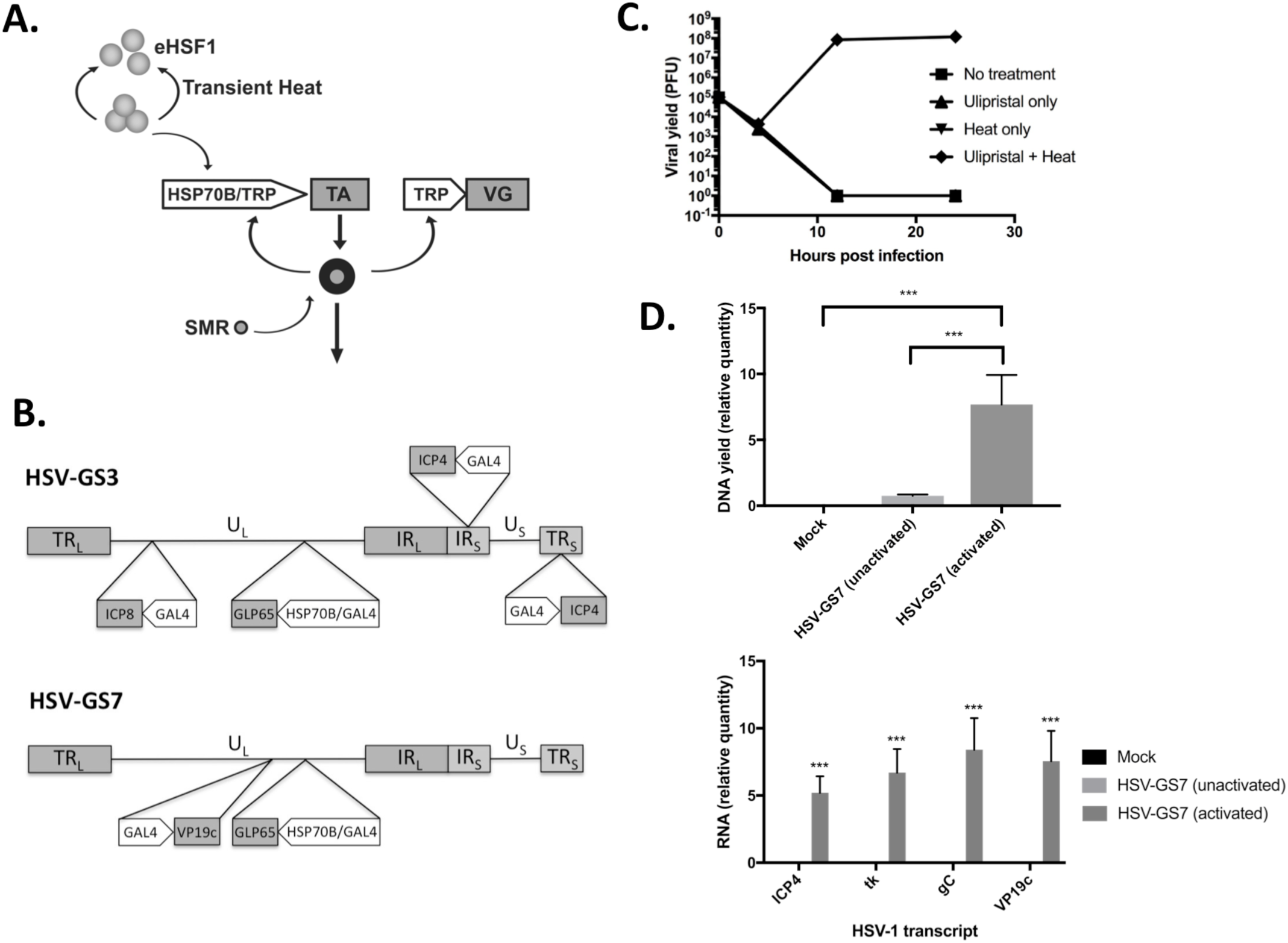
Two-component HSP70B promoter-based gene switch and replication-competent, controlled HSV-1-derived viral vectors. (A) Illustration of the operation of the dual-responsive gene switch employed to control replication of the viral vectors of the present study. (Copyright © American Society for Microbiology, (34)) eHSF1: endogenous heat shock transcription factor; HSP70B: HSP70B/HSPA7 promoter; TRP: transactivator-responsive promoter; TA: transactivator; VG: replication-essential viral gene; SMR: small-molecule regulator. (B) Diagrams of viral vectors HSV-GS3 (top of panel) and HSV-GS7 (bottom of panel). Both vectors contain, inserted into the intragenic region between the HSV-1 UL43 and UL44 genes, a GLP65 transactivator gene functionally linked to a promoter cassette consisting of a human HSP70B promoter and a GAL4-responsive promoter. In HSV-GS3, the promoters of the replication-essential genes encoding ICP4 and ICP8 were replaced with GAL4-responsive promoters. In HSV-GS7, the promoter of the replication-essential gene for VP19c was replaced with a GAL4-responsive promoter. TR_L_, TR_S_: long and short terminal repeats; U_L_, U_S_: long and short unique regions; IRL, IRS: long and short internal repeats. (C) Single step growth experiment for HSV-GS7 in Vero cells. Heat: exposure to 43.5°C for 30 min immediately after infection (i.e., immediately after removal of the viral inoculum). Ulipristal: 10 nM ulipristal was added to the medium at the time of initial infection. Each data point represents the mean of three individual assays. Error bars are not visible due to their range. pfu/ml values are shown. For details see under Materials and Methods. (D) Replication of HSV-GS7 (top graph) and expression of viral genes (bottom graph) in rear feet of mice (group size (n) of 5) determined 24 h after administration of the recombinant vector (50,000 pfu) to the plantar surfaces of the rear feet. HSV-GS7 replication was either not activated or was activated three h after infection by immersion of the infected rear feet in a 45°C water bath for 10 min in the presence of ulipristal. Ulipristal (50 µg/kg) had been administered IP at the time of initial infection. Twenty-four h after infection, animals were euthanized, and DNA and RNA were isolated from the infected rear feet. Levels of viral DNA were assessed by qPCR, and levels of ICP4, TK, gC and VP19c transcripts by RT-qPCR. Relative values and standard deviations are shown. *** p ≤ 0.05 for the comparison of “HSV-GS7 activated” and “HSV-GS7 unactivated” or “mock”.

## RESULTS

### Replication-competent controlled HSV-1 vectors

The two-component HSP promoter-based gene switch employed in the vectors of the present study relied on transactivator GLP65 that is activated by a narrow class of antiprogestins including mifepristone and ulipristal (31-34). GLP65 is a chimeric protein consisting of a DNA-binding domain from yeast transcription factor GAL4, a truncated ligand-binding domain from a human progesterone receptor and a transcriptional activation domain from human NFkappaB protein P65. A DNA segment containing a promoter cassette consisting of a human HSP70B promoter and a GAL4-binding site-containing (GAL4-responsive) promoter and a functionally linked GLP65 gene was inserted in between the UL43 and UL44 genes of HSV-1 wildtype strain 17*syn*+. This intermediate vector was utilized for the construction of vectors HSV-GS3 and HSV-GS7. In vector HSV-GS3, replication-essential immediate early gene RS1/ICP4 and early gene UL29/ICP8 had been placed under gene switch control by replacing the resident ICP4 and ICP8 promoters in the latter intermediate vector with GAL4-responsive promoters (Fig. 1B). In vector HSV-GS7, the replication-essential late gene UL38/VP19c had been subjected to gene switch control by substitution of its promoter with a GAL4-responsive promoter.

Replication of HSV-GS3 had previously been shown to be strictly dependent on activation by heat and antiprogestin in single-step growth experiments (34). When activated, the recombinant replicated nearly as efficiently as wildtype virus 17*syn*+. When administered to footpads of mice, HSV-GS3 only replicated subsequent to an activating treatment by locally administered heat in the presence of systemic antiprogestin. The results of a single-step growth experiment with HSV-GS7 are shown in Fig. 1C. The recombinant replicated similarly well as HSV-GS3 following heat treatment in the presence of antiprogestin ulipristal. No replication was observed after either heat or antiprogestin treatment, or in the absence of any treatment. To verify that HSV-GS7 replication was also tightly controlled *in vivo*, two of three groups of mice were administered HSV-GS7 virus (50,000 pfu per mouse) to the footpad, and the mice of one of the latter groups were given ulipristal intraperitoneally (i,p.) as well as, 3 h later, were subjected to a heat treatment to the footpads at 45°C for 10 min. One day later, all mice were euthanized, and DNA and RNA were extracted from their feet and analyzed by qPCR and RT-qPCR, respectively. Substantially larger amounts of HSV-1 DNA were detected in the feet of heat/ulipristal-treated mice than in not-treated mice (Fig. 1D). Expression of several viral genes was observed for activated virus but not for unactivated virus, strongly suggesting that viral replication and gene expression only occurred subsequent to heat/ulipristal activation.

It is noted that stocks of the recombinant viruses were prepared in cells that were subjected to daily heat treatment in the presence of antiprogestin until maximal cytopathic effect was reached. Titrations of the viruses were carried out on cells that provided missing proteins in *trans*.

### Protective immunity induced by activated HSV-GS3

Induction of protective immunity was evaluated in a mouse footpad lethal challenge model (35). In a first experiment, virus vectors were administered under anesthesia to the plantar surfaces of both rear feet of adult Swiss Webster outbred female mice (50,000 pfu per animal; 20 animals per group). Concurrently, and again 24 h later, the animals of one group received an intraperitoneal injection of 0.5 mg/kg of mifepristone. Three hours after inoculation, the mice of the latter group were subjected to heat treatment (43.5°C for 30 min) by immersion of their hindfeet in a temperature-controlled water bath. Twenty-two days later, all animals were challenged by a 20-fold lethal dose of HSV-1 wildtype strain 17*syn*+ administered by the same route as the original virus inoculum. Survival of the animals was followed until no more lethal endpoints were reached, i.e., until all surviving animals had fully recovered (Fig. 2). ICP4(-) replication-incompetent HSV-1 recombinant KD6 (36) induced a modest level of immunity. As had been expected, because it did not replicate and also did not express the major transcriptional regulator ICP4, unactivated HSV-GS3 provided a comparable degree of protection. Activated HSV-GS3 produced a far greater protective effect. Therefore, one round of efficient replication and/or expression of master regulator ICP4 resulting in expression of all viral proteins produced a dramatic enhancement of protective immunity. It is noted that the heat and antiprogestin treatment had no systematic effect on the level of protection induced by replication-defective virus KD6 (see under “Statistical evaluation of data from all comparable experiments”).

**FIG 2.**
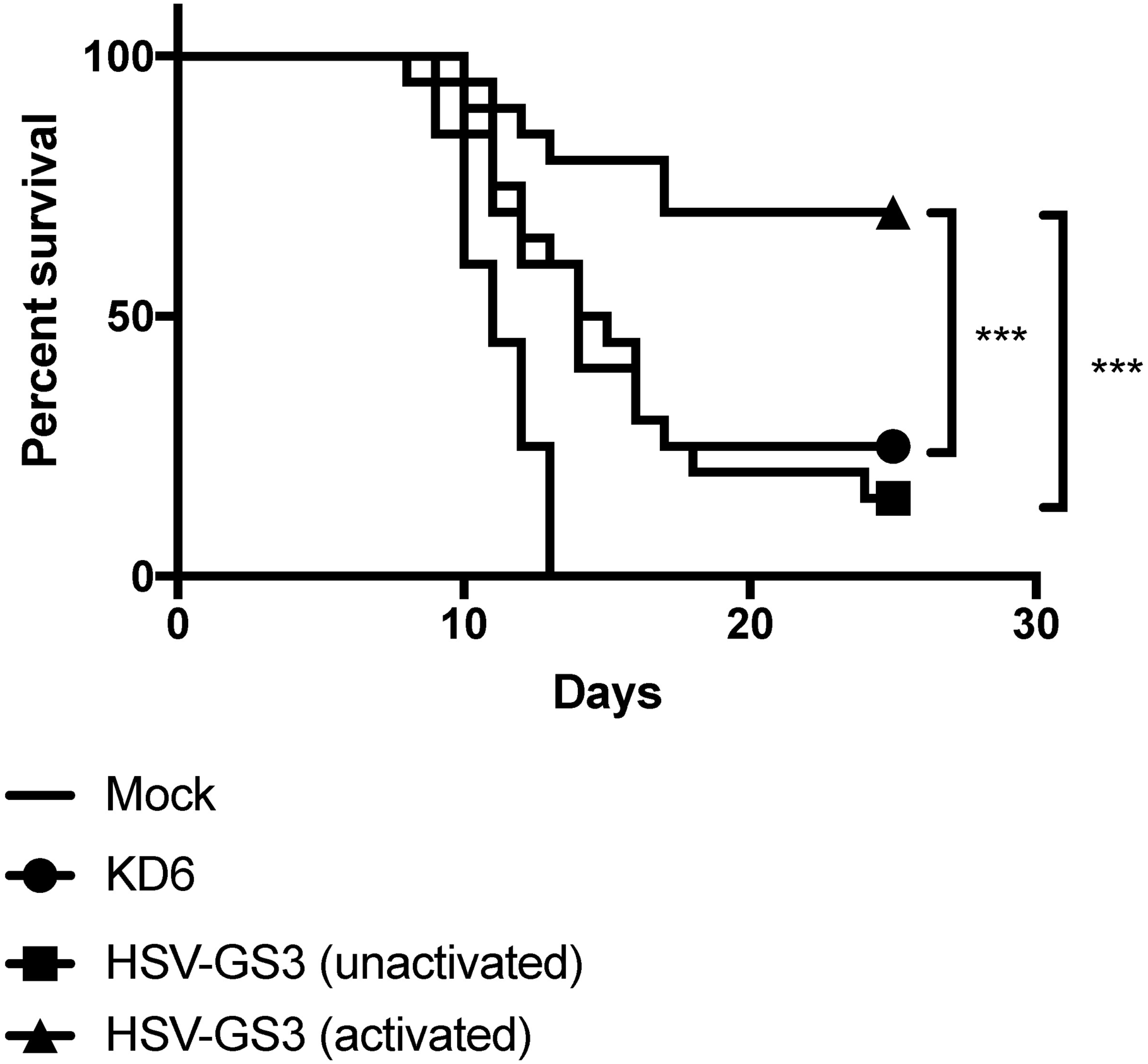
Immunization with HSV-GS3 protects against lethal challenge with HSV-1 17*syn*+ in the mouse footpad model. Groups of mice were inoculated on the plantar surfaces of both rear feet with 50,000 pfu/mouse of either the ICP4(-) HSV-1 replication-incompetent recombinant KD6 or the HSV-1 replication-competent controlled recombinant HSV-GS3 (two groups), or were mock-immunized with saline. Animals of one of the two HSV-GS3 groups received a 43.5°C/30min heat treatment 3 h after virus inoculation in the systemic presence of mifepristone (0.5 mg/kg administered IP at the time of inoculation) (“HSV-GS3 activated”), whereas animals of the other group were left untreated (“HSV-GS3 unactivated”). At 22 days post immunization, all mice were challenged with a 20-fold lethal dose of wild type HSV-1 strain 17*syn*+ applied to both rear feet. Data are presented as percent survival for each treatment group (n=20 for each treatment). *** p ≤ 0.05. See Materials and Methods for further details, and Table 2 and Fig. 6 for comparable experiments.

To determine whether immunization with activated HSV-GS3 reduced replication of the challenge virus more effectively than unactivated HSV-GS3 or replication-defective KD6 virus, additional groups of animals (5 animals per group) were immunized and challenged by wildtype virus as described above. Four days after challenge, the animals were euthanized, feet were dissected and homogenized, and virus present in the homogenates was titrated. Results revealed that activated HSV-GS3 virus reduced challenge virus replication by nearly two orders of magnitude (Table 1). Unactivated recombinant virus or KD6 were far less effective.

**TABLE 1.** Infectious virus (pfu) in the feet of mice 4 days post challenge

| Mock | KD6 | HSV-GS3<br>(unactivated) | HSV-GS3<br>(activated) |
| --- | --- | --- | --- |
| $5.5 \times 10^5 \pm 1.2 \times 10^4$ | $4.0 \times 10^4 \pm 8.3 \times 10^2$ | $9.4 \times 10^4 \pm 3.1 \times 10^4$ | $6.2 \times 10^3 \pm 1.5 \times 10^2$ |

We next examined in a similar experiment whether induction of enhanced protective immunity was caused by an activation of the heat- and antiprogestin-dependent gene switch that controls the expression of ICP4 and ICP8 in the HSV-GS3 recombinant. In this and all subsequent experiments, ulipristal was utilized instead of mifepristone, because the former compound is a more active gene switch co-activator than the latter compound (34). Furthermore, the heat treatment regimen was changed to heat exposure at 45°C for 10 min to achieve greater reproducibility between experiments. Results revealed that enhanced protection was only achieved following activation of HSV-GS3 by heat treatment in the presence of ulipristal (Fig. 3). Heat treatment alone or ulipristal alone did not significantly increase the protective effect over that produced by the unactivated virus. These findings strongly suggested that it was the activation of the dual-responsive gene switch in HSV-GS3 which prompted the expression of the regulated replication-essential genes as well as efficient virus replication that in turn resulted in the observed enhanced protective response. It is noted that in the latter experiment the virus inoculum was 500,000 pfu rather than 50,000 pfu as in most other experiments. This may account for the somewhat elevated level of protection induced by unactivated HSV-GS3.

**FIG 3.**
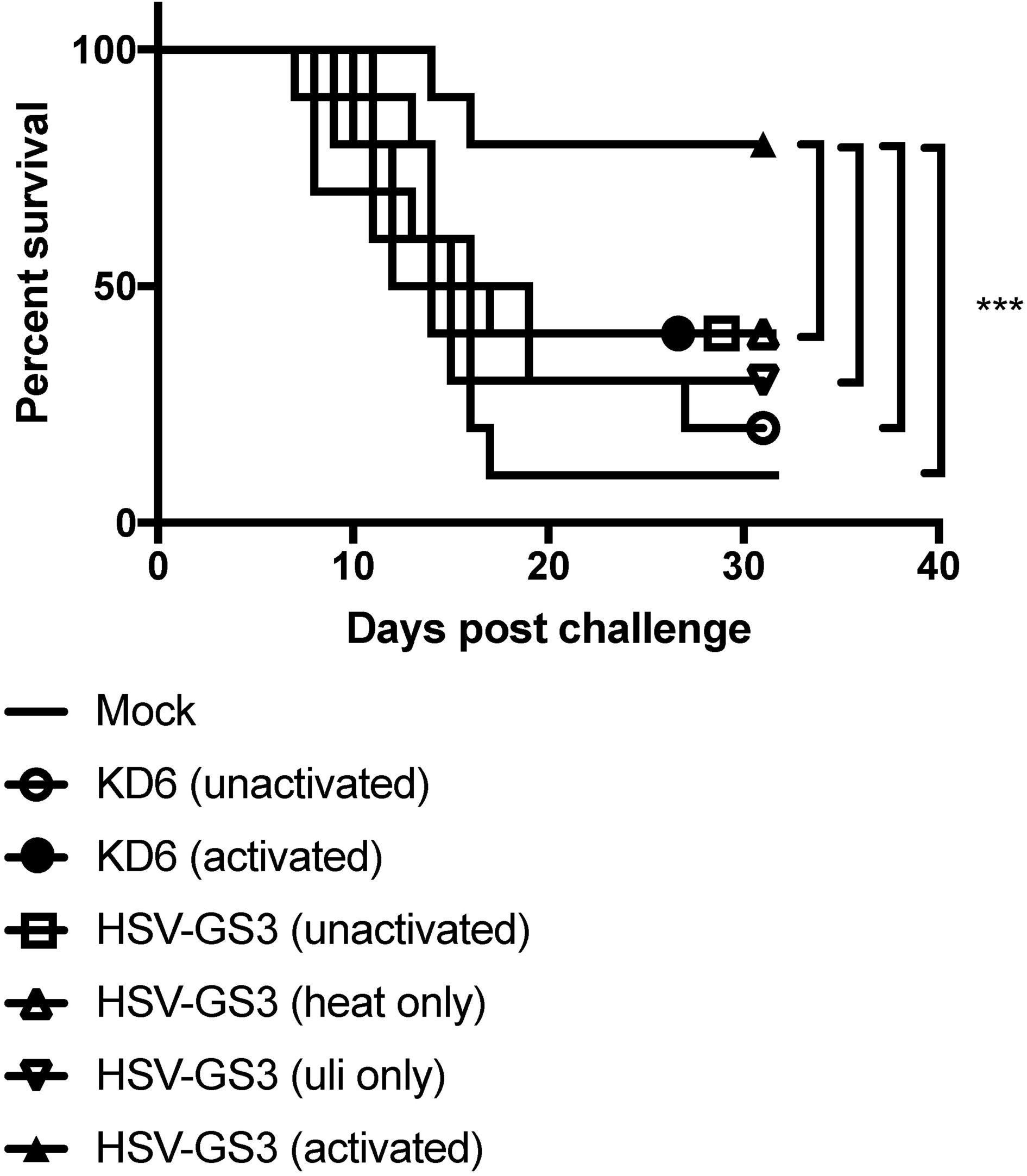
Gene switch-controlled immunization by recombinant HSV-GS3. Mice in four groups (n=10) were administered on both rear feet 500,000 pfu/mouse of HSV-GS3. The first HSV-GS3 group received a 45°C/10min heat treatment 3 h after inoculation in the systemic presence of ulipristal (50 µg/kg administered IP at the time of inoculation) (“HSV-GS3 (activated)”), the second group only received heat treatment (“HSV-GS3 (heat only)”), the third group was only administered ulipristal (“HSV-GS3 (uli only)”), and the fourth group was left untreated (“HSV-GS3 unactivated”). A further group was mock-immunized with saline. At 21 days, animals in the HSV-GS3 groups were boosted with another 500,000 pfu/mouse dose of HSV-GS3, and 10 days later all mice were challenged with a 20-fold lethal dose of wild-type HSV-1 strain 17*syn*+ applied to both rear feet. The data are presented as percent survival for each treatment group. *** p ≤ 0.05. See Materials and Methods for further details. Comparable data were obtained from a second experiment (not shown), in which mice were immunized with 50,000 pfu/mouse of HSV-GS3 and were not boosted.

### HSV-GS3 and HSV-GS7 induce comparable protective responses

In recombinant HSV-GS3, immediate early gene ICP4 and early gene ICP8 are subject to dual-responsive gene switch control. The greatly enhanced protective response that resulted from transient activation of the gene switch in HSV-GS3-immunized animals could have been due to the efficient virus replication triggered by the activation and/or to the activation of the genes for major transcription regulator ICP4 that in turn stimulated expression of all other viral genes in primarily infected cells. Another issue related to viral gene expression in cells that were secondarily (nonproductively) infected by HSV-GS3. Since the ICP4 genes are regulated in the recombinant virus, little expression of viral genes was expected to occur in the absence of another activating treatment. Abundant viral gene expression in secondarily infected cells (that may include antigen-presenting cells) conceivably may further enhance immune responses. Recombinant HSV-GS7 in which the late VP19c gene had been subjected to regulation had been constructed to examine these issues. HSV-GS3 and HSV-GS7 were compared in the mouse footpad challenge model. Results revealed that activated HSV-GS3 and HSV-GS7 provided similar levels of protection (Fig. 4). In the absence of activation, the protective effect of either recombinant was modest. Taking data from other experiments into account, the average protective effect for unactivated HSV-GS3 was 22% and that for unactivated HSV-GS7 was 13% (see under “Statistical evaluation of data from all comparable experiments”). The fact that the two recombinants which lacked expression of different proteins in the absence of activation induced comparable (modest) protective responses strongly suggested that it was not the absence of one or the other viral protein that limited the immune response to the recombinants. It was indeed the vigorous replication of the activated recombinants that enhanced the protective responses against them. In secondarily infected cells (in which HSF1 is not activated), HSV-GS7 was expected to express all viral proteins except the regulated capsid protein at approximately normal levels. In contrast, HSV-GS3 all but lacks ICP4 expression in the absence of activation and exhibits dramatically curtailed expression of other viral proteins (34). The finding that HSV-GS3 and HSV-GS7 induced comparable protective responses suggested that viral proteins expressed in secondarily infected cells did not contribute in a relevant fashion to the immune response. However, we cannot exclude the possibility that the immune system was sufficiently sensitive to detect the very low levels of viral antigen expression occurring in cells secondarily infected with HSV-GS3.

**FIG 4.**
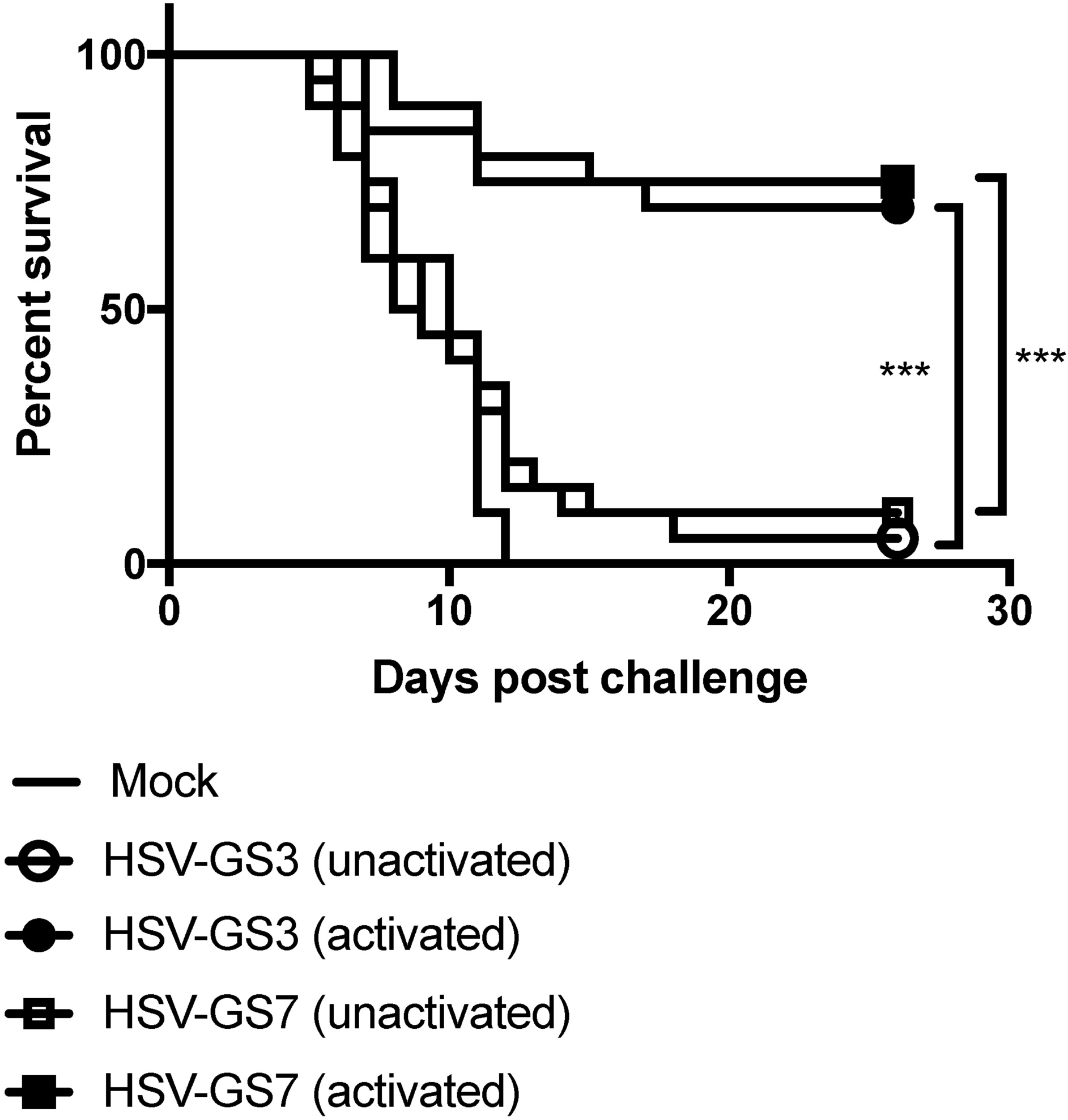
Immunization with HSV-1 replication-competent controlled recombinant HSV-GS7 and comparison with HSV-GS3. Mice were inoculated on both rear feet with 50,000 pfu/mouse of either HSV-GS3 or HSV-GS7 or were mock-immunized with saline. For each recombinant, one group of animals was subjected to heat treatment in the presence of ulipristal as described under Fig. 3 (“activated”), and another group did not receive this treatment (“unactivated”). At 21 days post inoculation, all mice were challenged with a 20-fold lethal dose of wild-type HSV-1 strain 17*syn*+ applied to both rear feet. The data are presented as percent survival for each treatment group (n = 20 for each HSV-GS3 or HSV-GS7 group; n = 10 for mock; *** p ≤ 0.05). See Table 2 and Fig. 6 for comparable experiments.

**TABLE 2.**
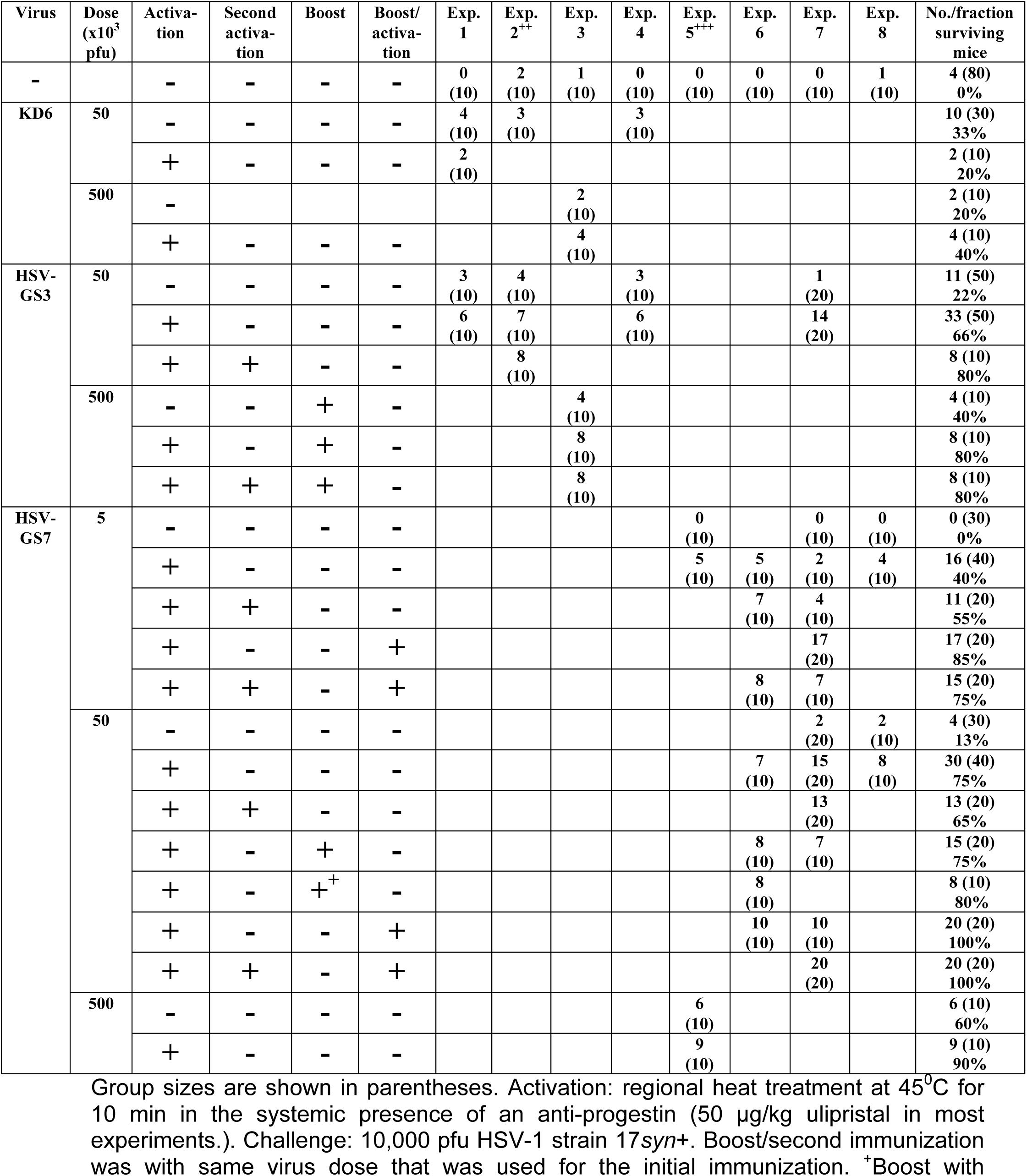
Number/fraction of survivors in immunization/challenge experiments.

### Second immunization further enhances protective immunity

We next investigated whether a second activation treatment applied two days after the first treatment (i.e., at a time at which the initial round of replication of HSV-GS7 should have been completed), or a second immunization would further enhance protection against wildtype virus challenge. The second immunization was administered three weeks after the initial virus application. Booster virus was either activated or left unactivated. In the part of an experiment reported on in Fig. 5A, groups of mice were immunized with 50,000 pfu/animal of recombinant HSV-GS7. Activation of the virus induced an immune response that protected 75% of animals after challenge with wildtype virus. After second immunization with a similar dose of HSV-GS7, all animals were protected. However, this only occurred if the booster virus was also subjected to an activating treatment. Second activation had no apparent effect. As effects of re-immunization may be more apparent in mice immunized with lower quantities of virus, the experiment also featured parallel groups of animals that were immunized and boosted with 5,000 pfu/animal of recombinant HSV-GS7. As Fig. 5B documents, second immunization followed by virus activation had an even more pronounced effect at the 10-fold reduced virus doses. It is noted that in this part of the experiment second activation appeared to modestly enhance the protective immune response.

**FIG 5.**
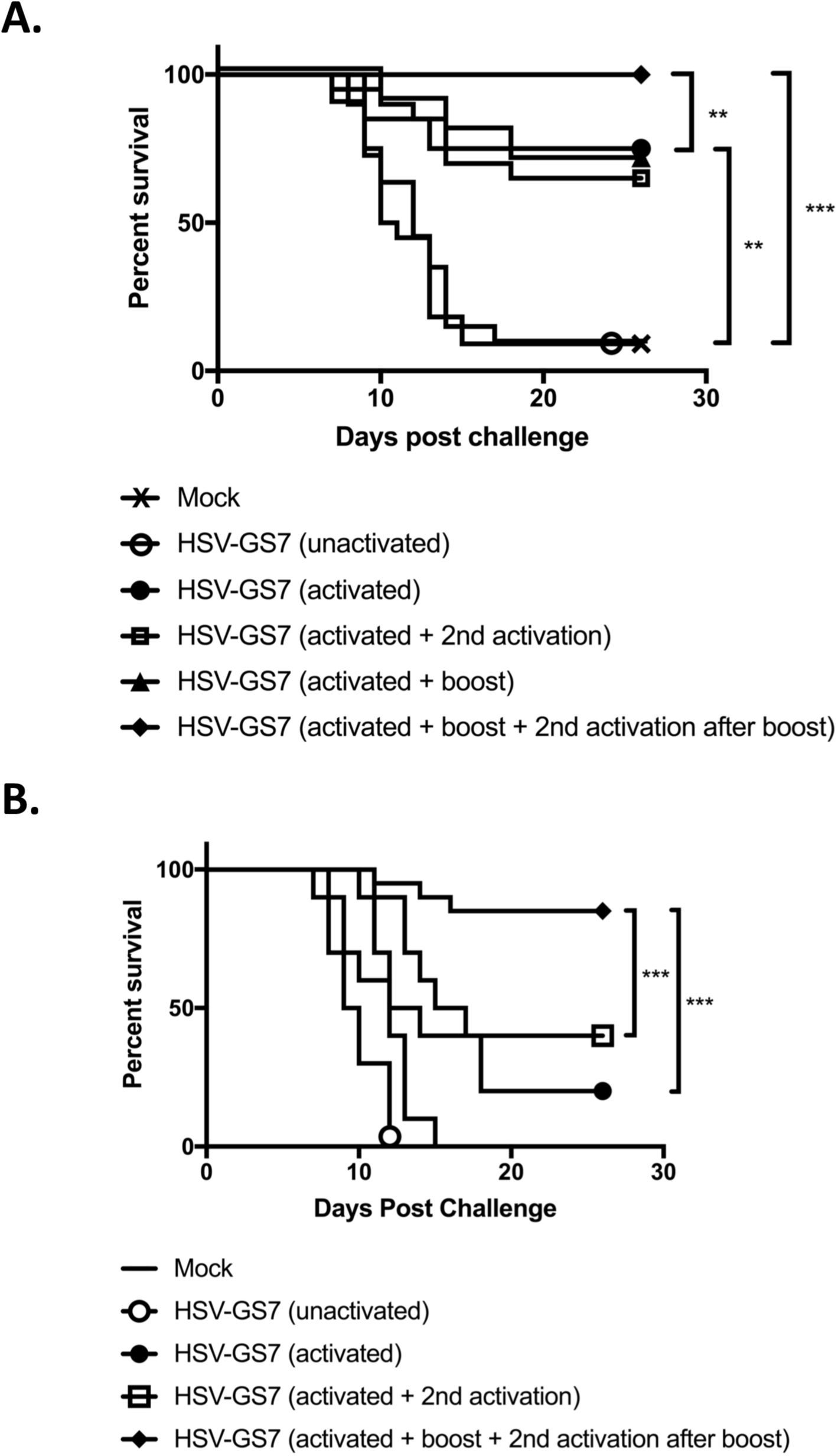
Effects of second immunization and second activation on HSV-GS7 efficacy. (A) Mice were inoculated on both rear feet with 50,000 pfu/mouse of HSV-GS7 or saline (mock; n = 10). HSV-GS7 groups: untreated (“unactivated”; n = 20), activated on day 1 (3 h after virus inoculation) by heat treatment at 45°C for 10 min in the presence of ulipristal (50 µg/kg administered IP at the time of inoculation) (“activated”; n = 20), activated on day 1 and reactivated 2 days later (“activated + 2nd activation”; n = 20), activated on day 1 and readministered 50,000 pfu/mouse of HSV-GS7 21 days later (“activated + boost”; n = 10), and activated on day 1, readministered 50,000 pfu/mouse of HSV-GS7 21 days after the first inoculation and reactivated 3 h later (“activated + boost + 2nd activation”; n = 10). Twenty-one days after the last treatment, all mice were challenged with a 20-fold lethal dose of wild-type HSV-1 strain 17*syn*+ applied to both rear feet. ** p ≤ 0.05; *** p < 0.01. (B) A similar experiment except that both initial and second immunizations were with 5,000 pfu/mouse of HSV-GS7 (n = 10; for second immunization n = 20; *** p ≤ 0.05). The data are presented as percent survival for each treatment group. See Table 2 and Fig. 6 for comparable experiments.

### Statistical evaluation of data from all comparable experiments

Relevant data from all immunization/challenge experiments that were conducted using essentially identical protocols are assembled in Table 2. Results of a meta-analysis of the HSV-GS7 data are presented graphically in Fig. 6. The difference in immunization efficacy between activated and unactivated HSV-GS7 was found to be highly significant. Also significant was the increase in protection afforded by second immunization with an activated HSV-GS7 vector. The effect of second activation two days after virus administration and first activation was not statistically relevant. Second activation tended to modestly enhance protective immunity in a majority of experiments in which this was addressed. However, in some experiments, e.g., in the experiment reported in Fig. 5A, an effect was not apparent. It is noted that the conditions for second activation had not been optimized.

**FIG 6.**
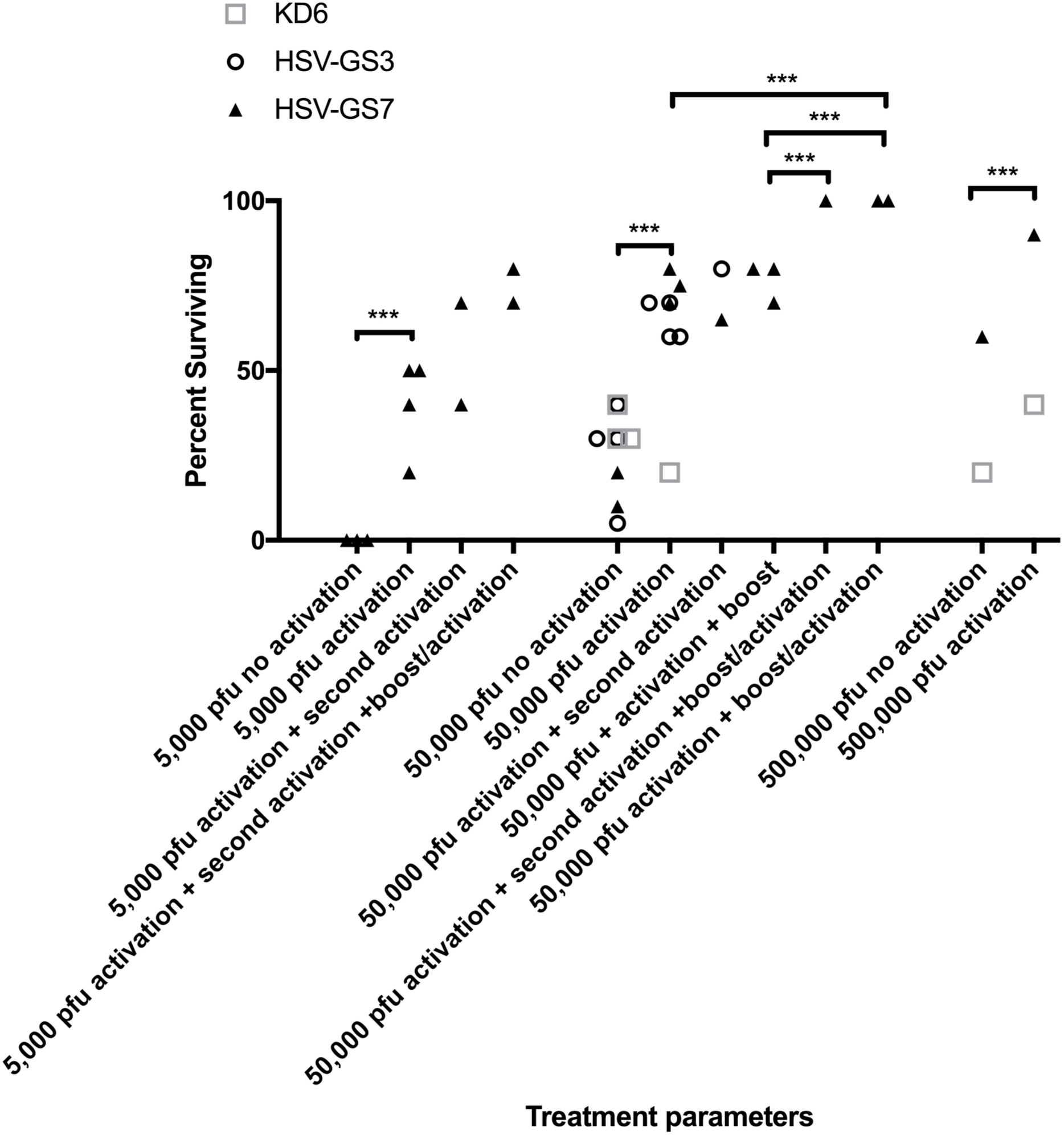
Meta-analysis of efficacy of HSV-GS7 in protecting mice against lethal HSV-1 challenge across comparable experiments. Selected data from Table 2 showing percent survival for different treatment conditions examined for KD6, HSV-GS3 and/or HSV-GS7 are graphically depicted. Data sets for HSV-GS7 were analyzed using a two-way ANOVA analysis. *** p ≤ 0.05.

### Immune correlates of protection against challenge

The presence and levels of antibodies capable of neutralizing HSV-1 strain 17*syn*+ were assessed in sera of mice that had been treated with activated or unactivated HSV-GS vectors, or with replication-incompetent virus KD6 (all at 50,000 pfu per animal) three weeks earlier. As expected, neutralizing antibodies could be detected subsequent to immunization with KD6 or unactivated HSV-GS viruses (Fig. 7A). Antibody levels in sera from KD6-immunized and unactivated HSV-GS vector-immunized animals were comparable. Both activated HSV-GS3 and HSV-GS7 induced clearly superior neutralizing antibody responses when compared with unactivated HSV-GS3 or HSV-GS7, or KD6. To assess cellular immune responses, HSV-1-specific T cells present in peripheral blood mononuclear cells (PBMCs) of mice that had been immunized three weeks prior were quantified by a modified limiting dilution lymphoproliferation assay. Both unactivated HSV-GS vectors and KD6 virus were found capable of inducing detectable levels of HSV-1-specific T cells (Fig. 7B). Substantially better responses were produced by activated HSV-GS3 and HSV-GS7.

**FIG 7.**
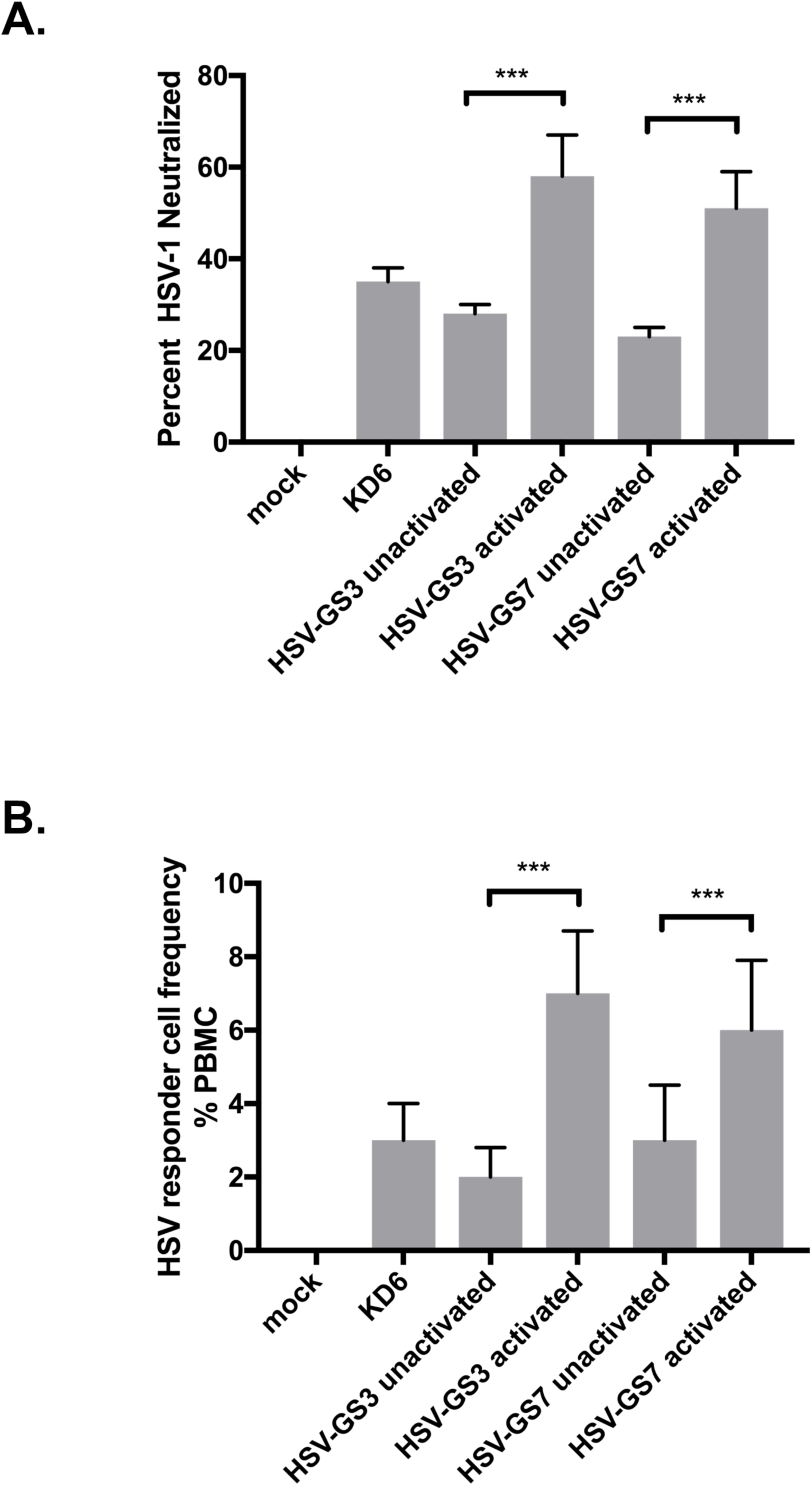
Anti-HSV-1 humoral and cellular immune responses. (A) Neutralizing antibodies induced by immunization. Dilutions of mouse serum samples (n = 5 for each experimental group) were tested for their ability to neutralize HSV-1 as described in Materials and Methods. Values are presented as averaged percentages of HSV-1 pfu neutralized for each group. ANOVA: *** p ≤ 0.05. See Materials and Methods for experimental detail. (B) Responder cell frequency. The number of HSV-1-specific lymphocytes induced three weeks after immunization was determined by a limiting dilution lymphocyte proliferation assay. The data are presented as averaged responder cell frequency for each experimental group. Three blood/PBMC samples were analyzed per group. ANOVA: *** p ≤ 0.05. See Materials and Methods for experimental detail.

### Immune responses to a heterologous antigen presented in the context of vigorous viral replication

To evaluate immune responses to a heterologous antigen expressed from a HSV-GS vector, recombinant HSV-GS11 was constructed. The recombinant was derived from HSV-GS3 by the insertion in the UL37/UL38 intergenic region of an expression cassette comprising a gene for an equine influenza hemagglutinin (EIV Prague/56 HA) functionally linked to a CMV (immediate early) promoter (Fig. 8A). Expression of the influenza gene was verified in groups of adult female BALB/c mice inoculated on the rear footpads with either saline, HSV-GS3 or HSV-GS11 (50,000 pfu per animal). The HSV-GS3 group as well as one of the HSV-GS7 groups were subjected to activation treatment. Another HSV-GS7 group underwent an activation treatment twice, the second treatment administered two days after the first treatment. One day after the last treatment, animals were euthanized, and RNA was extracted from one hindfoot and protein from the other. Results from an RT-qPCR analysis of HA gene expression are shown in Fig. 8B, and results from an EIV Prague/56 HA-specific ELISA in Fig 8C. HA RNA and HA could be detected in samples from HSV-GS11-inoculated animals, most abundantly when the animals had been subjected to activation treatment. Both HA RNA and protein levels appeared to be somewhat higher in twice-activated animals than in once-activated animals.

**FIG 8.**
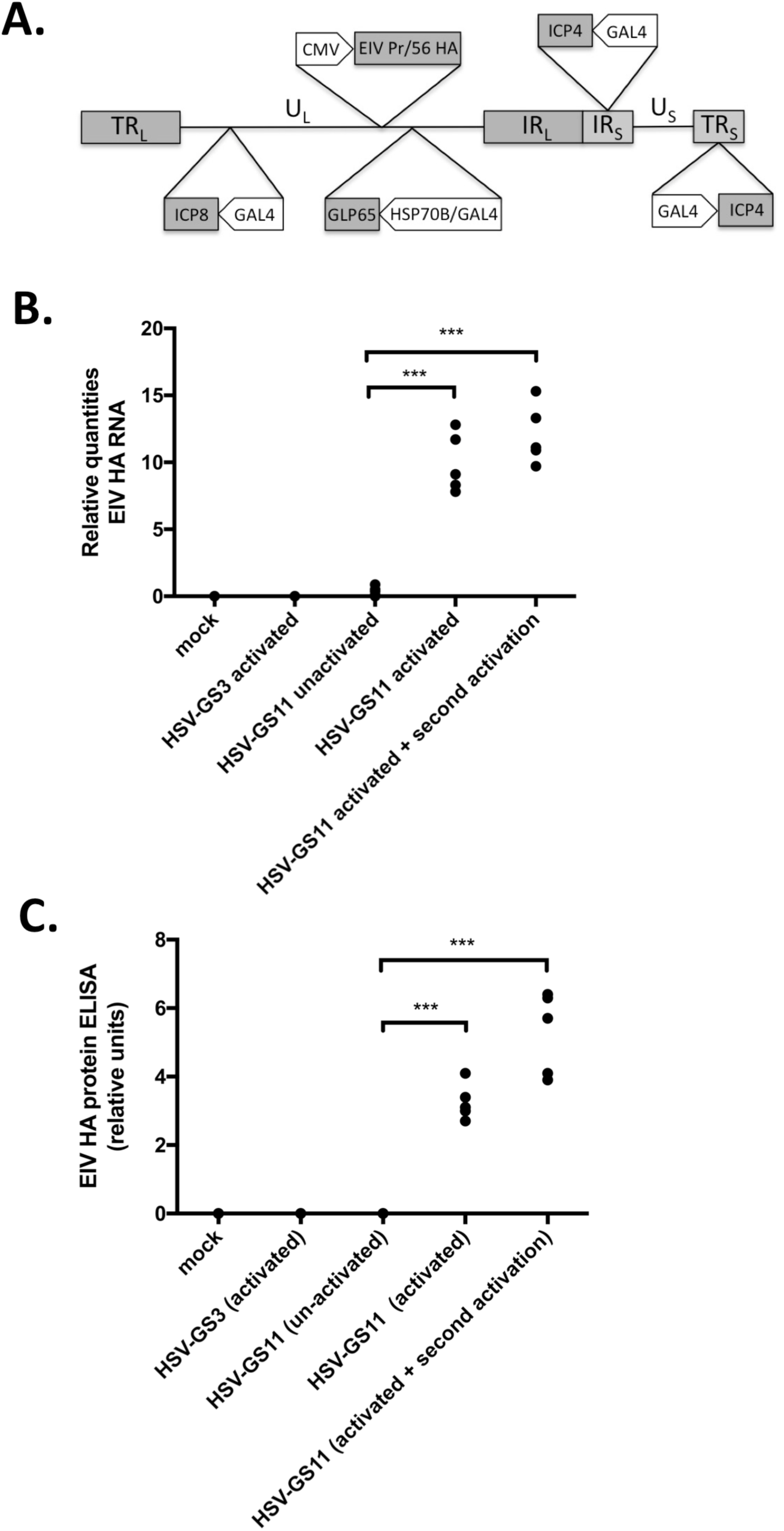
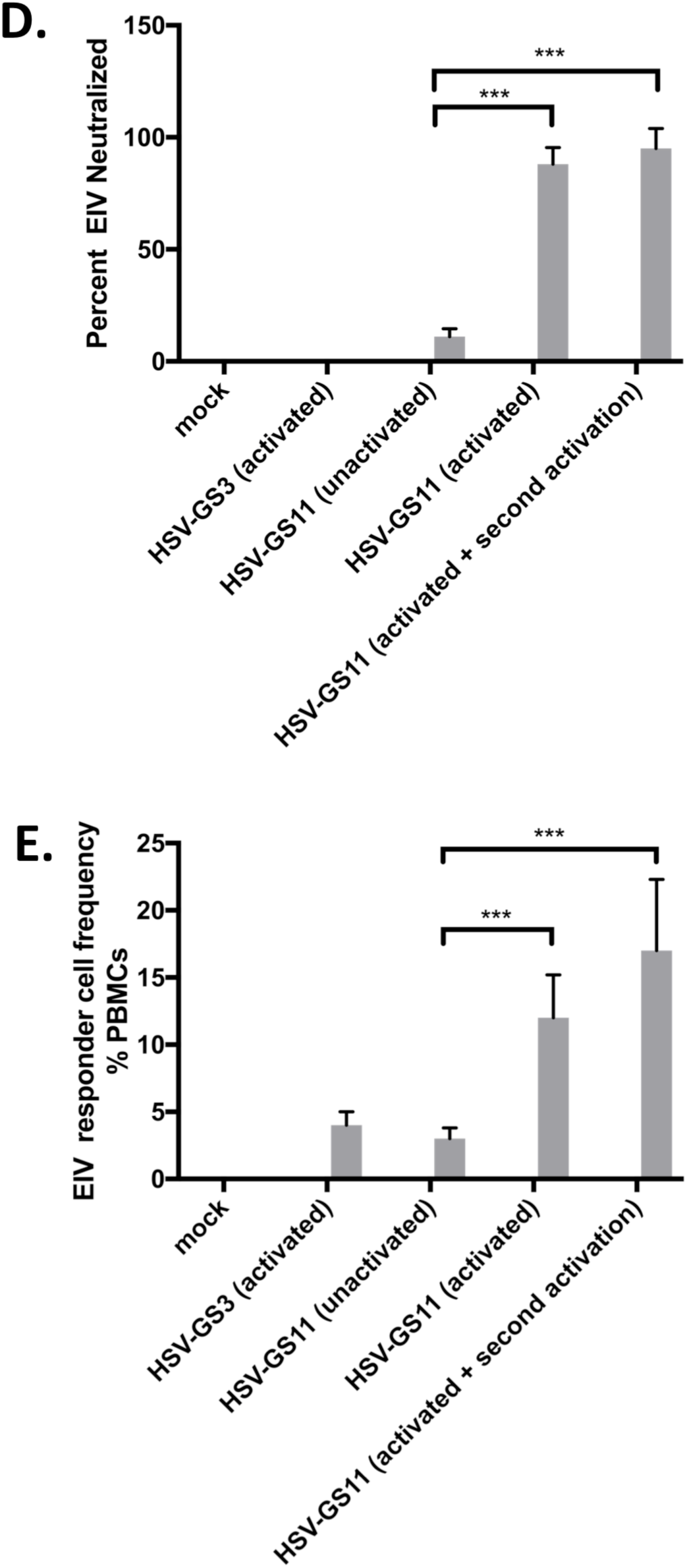
Immunization with HSV-GS11 induces a robust immune response against EIV Prague/56 HA. (A) HSV-GS11 vector containing the EIV Prague/56 (H7N7) HA gene driven by the CMV IE promoter and inserted into the intergenic region between UL37 and UL38. The HSP70/GAL4-GLP65 TA cassette is inserted into the intergenic region between UL43/44. The promoters of the essential HSV-1 genes ICP4 and ICP8 are replaced with GAL4-responsive promoters. (B) Detection of EIV HA RNA expression following immunization of mice with HSV-GS11. Groups (n = 5) of 6-8-week old female BALB/c mice were inoculated on both rear footpads with either saline (mock), HSV-GS3 (50,000 pfu) or HSV-GS11 ((50,000 pfu) as described under Materials and Methods. Vector replication was activated in some treatment groups by administration of heat and ulipristal as described in Fig. 3. One treatment group received a second activation treatment that was administered two days after the first activation treatment. Mice feet were harvested 24 hours after the last treatment, and RNA was isolated and cDNA prepared. Samples were analyzed by qPCR with EIV Prague/56 HA-specific Taqman primers/probe (Table 3). Data are presented in relative units of fluorescence. ANOVA: *** = p ≤ 0.05. (C) Detection of EIV HA antigen by ELISA following immunization of mice with HSV-GS11. Groups (n = 5) of 6-8-week old female BALB/c mice were inoculated on both rear footpads with either saline (mock), HSV-GS3 (50,000 pfu) or HSV-GS11 ((50,000 pfu) and were subjected to the same treatments as in B. Mice feet were harvested 24 hours after the last treatment, and protein homogenates were analyzed using an EIV Prague/56 HA-specific ELISA as described under Materials and Methods. Data are presented in relative units of fluorescence. ANOVA: *** = p ≤ 0.05. (D) Neutralizing antibodies induced by immunization. Groups (n = 5) of 6-8-week old female BALB/c mice were administered HSV-GS3, HSV-GS7 or saline (mock) and were treated as in B. Twenty-one days post immunization, mouse serum samples were tested for their ability to neutralize EIV Prague/56. Values are presented as percent of EIV Prague/56 HA pfu neutralized. (E) Responder cell frequency assay. Immunizations and treatments were as in D. Twenty-one days post immunization, the number of EIV Prague/56 HA-specific lymphocytes was determined by a limiting dilution lymphocyte proliferation assay. The data are presented as responder cell frequency of each experimental group.

**TABLE 3.**
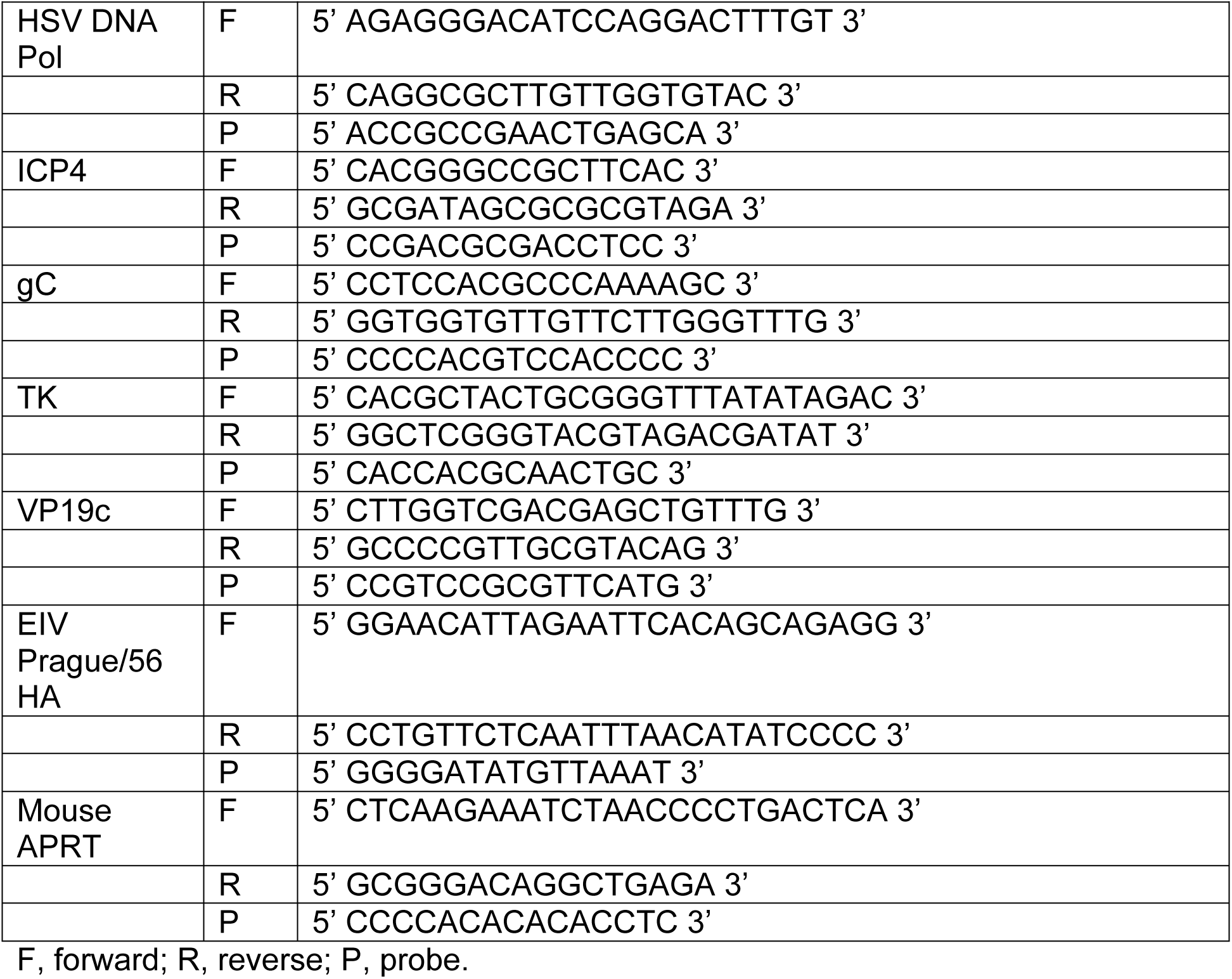
Primers and probes used for qPCR.

To assess immune responses (three weeks after immunization), additional groups of mice were inoculated with saline (mock immunization), or with 50,000 pfu of HSV-GS3 or HSV-GS11 (three groups). As in the above-described part of the experiment, all animals in the HSV-GS3 group as well as in two of the HSV-GS11 groups were subjected to an activation treatment. The animals of one of the latter groups received a second activation treatment two days later. Serum samples were tested for their ability to neutralize EIV Prague/56. As expected, neutralizing antibodies were not detected in unimmunized (not shown), mock-immunized or vector-immunized animals (Fig. 8D). Unactivated HSV-GS11 was capable of inducing a neutralizing antibody response. Activation of HSV-GS11 shortly after inoculation resulted in a several-fold magnified response. It is noted that twice-activated HSV-GS11 elicited an only marginally better response than once-activated virus. HA-specific T cells present in PBMCs were quantified by the same type of responder cell frequency assay that had been used to assess numbers of HSV-1-specific T cells. HA-specific T cells were not detected in unimmunized (not shown) or mock-immunized animals (Fig. 8E). Induction of a T cell response was observed in animals immunized with HSV-GS11 but not subjected to an activation treatment. Activated vector (HSV-GS3) produced a similar response. Far greater numbers of HA-specific T cells were found in animals immunized with once or twice activated HSV-GS11.

## DISCUSSION

Once-activated replication-competent controlled HSV-1 recombinants HSV-GS3 and HSV-GS7 induced considerably more potent protective immune responses than replication-defective HSV-1 strain KD6. Twice immunized animals were completely protected against lethal challenge. Levels of partial protection engendered by unactivated HSV-GS3, unactivated HSV-GS7 and KD6 (lacking expression of different viral proteins) were comparable, which strongly suggests that the much-enhanced protection provided by the activated HSV-GS recombinants was a consequence of their ability to efficiently replicate. Both better neutralizing antibody and better T cell responses were induced by the activated HSV-GS vectors, indicating that the replicating vectors broadly enhanced immune responses and not only boosted cellular immunity. Generalizing these findings, it appears that replication-competent controlled vectors are promising new agents that may better protect against diseases caused by the viruses from which they were derived than replication-incompetent/inactivated vaccines and, possibly, also live attenuated vaccines. What is perhaps even more important, our observations suggest that replication-competent controlled vectors can be excellent immunization platforms that may be exploited for the elaboration of new “vaccines”. Expression of an influenza antigen induced far more potent neutralizing antibody and antigen-specific T cell responses when the antigen was expressed in the context of vector replication than in the absence of vector replication. To the extent that such a comparison is valid, neutralizing antibody responses as well as antigen-specific T cell responses against the influenza antigen elicited by activated HSV-GS11 were both stronger than the corresponding responses against vector antigens.

A second activation treatment administered two days after the initial activation of a replication-competent controlled HSV-1 vector did not consistently enhance the protective immune response against challenge by the wildtype virus from which the vector was derived. Also, humoral and cellular immune responses against an influenza virus antigen expressed from a replication-competent controlled HSV-1 vector were only marginally enhanced by a second activation treatment. At this time, we cannot distinguish the possibilities that the second activation was only marginally effective because the immune system was already maximally engaged by the initial immunization or because optimal conditions for the second activation treatment had not been identified. Additional research is required to resolve this question.

Immunization by replication-competent controlled vectors represents a novel paradigm that may be elaborated in various ways. Therefore, the work presented herein should be regarded as a first illustration of the principle. In general, any of a number of DNA viruses including members of all herpesvirus subfamilies may be employed for the construction of replication-competent controlled vectors. We selected a fully virulent HSV-1 strain as the backbone of the GS vectors for several reasons, including that HSV have a double-stranded DNA genome that does not integrate in the host genome and are tolerant of sizeable insertions, support stable expression of heterologous genes, replicate and lyse cells efficiently as well as respond to antiviral drugs (37-40). Although HSV are endemic in the human population, available data indicate that pre-existing immunity is not an obstacle to vaccination/immunization uses of HSV vectors (discussed in ref. 34). Furthermore, latent infection was considered a manageable issue because replication of replication-competent controlled vectors can be stringently controlled, and reactivation should not be capable of occurring in the absence of activation.

Regarding the two-component HSP promoter-based gene switch that is used for controlling viral replication, several ligand-dependent transactivators are known that may be considered for incorporation in such a gene switch. They include transactivators that are derived from the bacterial tetracycline repressor or other bacterial repressors as well as chimeric transactivators that exploit ligand-binding domains of various steroid receptors, e.g., human progesterone, human estrogen or insect ecdysone receptors, or are activated by dimerization (28). We previously constructed and analyzed two-component HSP promoter-based gene switches that are co-activated by antiprogestins, ecdysteroid ponasterone A and rapamycin/rapalog, respectively (31,41). For regulating the HSV-GS vectors, we made use of a heat- and antiprogestin-activated gene switch that is strictly dependent on both heat and antiprogestin. While it has been more thoroughly characterized than any other (31), this gene switch also recommends itself for several other reasons. It comprises transactivator GLP65 that is activated by a narrow spectrum of antiprogestins but not progestins (34). Since it only includes a truncated ligand-binding domain of a progesterone receptor, GLP65 should not be susceptible to ligand-independent activation; in fact, after many years of experimental use, no antiprogestin-independent activation has been reported. At the low doses at which they activate GLP65, antiprogestins such as mifepristone and ulipristal are not expected to have any negative effects on the immune system (26). The latter antiprogestins also have been safety-tested in human subjects as well as are readily available. They are not considered essential active ingredients by the WHO, and their primary use has been in emergency contraception. To maintain maximal safety in a human immunization scenario, i.e., to preclude any possibility of reactivation from latency, antiprogestins should not be administered subsequent to immunization with a HSV-GS-type vector. Therefore, the vectors may not be employed for immunization of healthy female subjects of reproductive age or younger, or the subjects may forego later use of antiprogestins for emergency contraception. In contrast, the HSV-GS vectors may be well suited as immunization agents or platforms in middle-aged or elderly subjects. It is noted that it may be feasible to disable replication of fully activated HSV-GS vectors in latently infected cells through further engineering.

Replication-competent controlled virus vectors may be capable of making a unique contribution to immunization of immunocompromised patients. A recent study estimated that in 2012 over 3% of the adult populations of the United States and the United Kingdom were immunocompromised (42). Frequencies in other upper-income countries are expected to be comparable. These patients are particularly vulnerable to vaccine-preventable disease and would greatly benefit from effective vaccination. Owing to their condition, their immune response to inactivated or subunit vaccines may be inadequate. Unfortunately, use of live attenuated vaccines that may produce a more balanced and more persistent immune response is problematic because of the perceived danger that such vaccines may be insufficiently controlled by the weakened immune system and may cause the disease they are intended to prevent. According to most relevant guidelines, live attenuated vaccines are contraindicated in patients under immunosuppressive therapy or HIV patients with low CD4 count (43). Hence, use of currently marketed vaccines against tuberculosis (BCG), measles, mumps and rubella, varicella-zoster, rotavirus, influenza (nasal) and yellow fever is universally discouraged. Newer more effective vaccines under development, e.g., new pertussis vaccine BPZE1 (44), also may not become available to the immunocompromised. Replication-competent controlled virus vectors expressing antigens of the latter microbes in the context of efficient viral replication can be expected to elicit similarly or more potent immune reactions as the contraindicated live attenuated vaccines. Escape from immune control will not be an issue because expression of replication-essential genes is under stringent dual control. Hence, replication of the vectors can be activated and safely disactivated independent of the immune status of a patient.

In this report, we have focused on immunization effects of once-activated replication-competent controlled vectors that undergo one round of replication. We note that viral vectors that were intended to be limited to one cycle of replication were advanced before as potential vaccines (45). These so-called disabled infectious single-cycle (DISC) viruses typically were deleted for the gene for glycoprotein H which is critical for infection. Juxtaposing DISC viruses and replication-competent controlled viruses, the latter viruses replicate nearly as efficiently as the wildtype virus that had been used for their construction. No comparable information is available for the DISC viruses that were advanced as vaccine candidates. Replication of DISC viruses is not localized but occurs throughout the host. Depending on the dose of virus administered, a single round of replication of disseminated virus can be expected to result in significant toxicity. Not surprisingly, intracranial administration of DISC virus TA-HSV resulted in neurotoxicity that was only about two orders of magnitude lower than that of the wildtype virus from which the strain was derived (46). Replication-competent controlled viruses should be devoid of such toxicity because they are virtually incapable of replicating unless deliberately activated. It is noted that the stringency of the replication block can be increased further by subjecting additional replication-essential genes to gene switch control. Furthermore, even though progeny virus produced by a DISC virus upon immunization is not supposed to be capable of re-infecting cells, latent infection has been observed (46). Latent infection is known to involve spread from cell to cell (47). Hence, as has been pointed out before (48), DISC viruses such as TA-HSV appear to be leaky for replication *in vivo*. The above properties of DISC viruses are not desirable in an immunization agent or vector administered to healthy people and would disqualify such viruses from use in persons with impaired immune systems. It is noted that a clinical trial was conducted that investigated a DISC virus as a therapeutic vaccine against recurrent genital herpes (49). Results were negative. The DISC virus (TA-HSV) studied had been derived from a minimally virulent HSV-2 wildtype strain (46,50). Moreover, the virus was administered at a relatively low dose, apparently to avoid toxicity. Failure of the trial should not be very surprising.

Unlike a conventional vaccine, a replication-competent controlled vector will need to be activated after administration to a subject to be immunized. Activation will need to be neither costly nor inconvenient. A single ulipristal dose can be administered orally at the time of immunization. Regarding heat treatment, we have developed small pads that conduction-heat the inoculation region by crystallization of a supercooled salt solution (51). The pads are capable of maintaining a temperature of 45°C for 15 min. This heat dose was shown to be sufficient for strong activation in all layers of the skin of human subjects of the HSP70B promoter, which is the promoter used in the two-component gene switch that controls the HSV-GS vectors.

## MATERIALS AND METHODS

### Cells, plasmids and viruses

Rabbit skin (RS) cells were a gift from E. Wagner and were propagated in minimal essential medium Eagle’s salts (MEM) (Life Technologies/Thermo Fisher Scientific) supplemented with 5% heat-inactivated calf serum (Atlanta Biologicals, Lawrenceville, GA), 292 mg/ml L-glutamine, 250 U/ml of penicillin and 250 µg/ml of streptomycin (Life Technologies). Vero cells (procured from the American Type Culture Collection (ATCC) and Vero-derived E5 cells (52) (obtained from N. DeLuca) were propagated in Dulbecco’s modified Eagle’s medium (DMEM) with 10% heat-inactivated fetal calf serum, 250 U/ml of penicillin and 250 µg/ml of streptomycin. HSV-1 strain 17*syn*+ and ICP4 deletion strain KD6 (36) were obtained from J. Stevens. It is noted that stocks of KD6, HSV-GS3 and HSV-GS7 were propagated on E5 cells. For HSV-GS3 and HSV-GS7, medium was supplemented with 10 nM mifepristone, and infected cultures were subjected to daily heat treatment at 43.5°C for 30 min for three successive days. EIV Prague/56 was a gift of T. Lengsfeld and was propagated on Madin-Darby canine kidney (MDCK) cells (ATCC) in DMEM supplemented with 10% heat-inactivated horse serum as previously described (53). Virus stocks of EIV were titrated using TCID50 assays in MDCK cells. All cells were cultured at 37°C under 5% CO_2_.

### Chemical reagents

Mifepristone was obtained from Sigma-Aldrich, and ulipristal acetate was from D-Innovation Pharmaceutical Inc., Chengdu, Sichuan, PR China. Both compounds were USP grade.

### Construction of viruses

All viruses were constructed using wild type HSV-1 strain 17*syn*+ as the backbone. Viral recombinants were generated by homologous recombination of engineered plasmids along with purified virion DNA in RS cells transfected by the calcium phosphate precipitation method as previously described (40). Plasmids used to engineer the insertions of the transactivator cassette or the GAL4-responsive promoters comprised HSV-1 sequences cloned from strain 17*syn*+. HSV-GS3 (Fig. 1B) was constructed as previously described (3,34). HSV-GS7 was derived from intermediate vector HSV-17GS43 (34) that contains an insertion between the UL43 and UL44 genes of a transactivator cassette containing a GLP65 gene under the control of a promoter cassette that combined a human HSP70B promoter and a GAL4-responsive promoter (31). To place the UL38/VP19c gene under regulation of a GAL4-responsive promoter, plasmid pBS-KS:GAL4-UL38 was constructed. This plasmid contains a GAL4 promoter inserted in between the HSV-1 UL38 recombination arms of plasmid pBS-KS:UL38∆promoter. A GAL4-responsive promoter comprising six copies of the yeast GAL4 UAS (upstream activating sequence), the adenovirus E1b TATA sequence and the synthetic intron Ivs8 was excised from plasmid pGene/v5-HisA (Invitrogen, now Life Technologies) with AatII and HindIII, and the resulting 473 bp fragment was gel-purified. For the vector, pBS-KS:UL38∆promoter was digested with AatII and HindIII, and the resulting 4285 bp fragment was gel-purified and shrimp alkaline phosphatase-treated. Ligation of the latter two fragments placed the GAL4 promoter in front of the UL38 transcriptional start site. To produce recombinant HSV-GS7, RS cells were co-transfected with plasmid pBS-KS:GAL4-UL38 and purified HSV-17GS43 virion DNA. Subsequent to the addition of mifepristone to the medium, the co-transfected cells were exposed to 43.5°C for 30 min and then incubated at 37°C. Subsequently, on days 2 and 3, the cells were again incubated at 43.5°C for 30 min and then returned to 37°C. Picking and amplification of plaques, screening and plaque purification was performed essentially as described for HSV-GS3 (3,34). The resulting plaque-purified HSV-GS7 was verified by Southern blot as well as by PCR and DNA sequence analysis of the recombination junctions. Plasmid pBS-KS:UL38∆promoter was constructed by deletion of the region from −1 to −47 of the UL38 promoter, i.e., by synthesizing two PCR fragments (one 437 bp and the other 550 bp) on either side of the deletion and cloning these into pBS KS+. HSV-GS11 was derived from the vector HSV-GS3 (34) and contains an insertion between the UL37 and UL38 genes of a gene cassette expressing the EIV Prague/56 HA gene driven by the CMV IE promoter. A recombination plasmid was constructed by the following sequential steps. First, a 814 bp fragment containing the region spanning the HSV-1 UL37/UL38 intergenic region from nt 83,603-84,417 from plasmid NK470 was subcloned into pBS that had had the MCS removed (digestion with KpnI/SacI) to yield pBS:UL37/38. A cassette containing a synthetic CMV IE promoter flanked by the pBS-SK+ MCS was ligated into pBS:UL37/38 digested with BspE1/AflII, which enzymes cut between the UL37 and UL38 genes, to yield the plasmid pIN:UL37/38. The EIV Prague/56 HA gene was PCR cloned from cDNA prepared from EIV Prague/56. Briefly, RNA was prepared by Trizol extraction of a stock of EIV Prague 56 and was reverse transcribed using Omni-Script Reverse Transcriptase (Qiagen) according to the manufacturer’s instructions. The cDNAs were cloned into pBS, and the clone containing the HA gene (pBS-EIVPrague56/HA) was confirmed by sequence analysis. The Prague/56 HA gene was subcloned from this plasmid and inserted behind the CMV promoter in the plasmid pIN:UL37/38 to yield plasmid pIN:37/38-Prague56/HA. To produce recombinant HSV-GS11, RS cells were co-transfected with plasmid pIN:37/38-Prague56/HA and purified HSV-GS3 virion DNA. Subsequent to the addition of mifepristone to the medium, the co-transfected cells were exposed to 43.5°C for 30 min and then incubated at 37°C. Subsequently, on days 2 and 3, the cells were again incubated at 43.5°C for 30 min and then returned to 37°C. Picking and amplification of plaques, screening and plaque purification was performed essentially as described for HSV-GS3 (3,34). The resulting plaque-purified HSV-GS11 was verified by Southern blot as well as by PCR and DNA sequence analysis of the recombination junctions.

### Construction of expression plasmid pVP19c

Plasmid pVP19c was constructed by PCR-cloning of the promoter and coding sequence of the HSV-1 VP19c gene into pBS KS+ using HSV-1 strain 17*syn*+ virion DNA as the PCR target.

### Single step growth analysis

Confluent monolayers of Vero cells were infected with HSV-GS7 at a multiplicity of infection (MOI) of 3. Virus was allowed to adsorb for 1 h at 37°C; then the inoculum was removed, and the cells were overlaid with complete medium. Ulipristal treatment (10 nM) was initiated at the time of the initial infection. Heat treatment was performed either immediately after infection or 4 h later by floating the sealed dishes in a 43.5°C water bath for 30 min. The dishes were then incubated for 72 h at 37°C. At 0, 4 or 8, 12 or 16, and 24 or 28 h post-infection, two dishes were removed, and the cells were scraped into medium for harvesting and subjected to two freeze-thaw cycles. Infectious virus was then determined by titrating the lysate of each dish in triplicate on 24 well plates of confluent E5 cells transfected 24 h prior to infection with expression vector pVP19c using Lipofectamine 2000 (Life Technologies). Plaques were visualized 2 days after infection using an antibody plaque assay, essentially as previously described (34).

### Immunization

Briefly, 4-6-week old female outbred Swiss Webster mice (Envigo, Tampa, FL) (or BALB/c mice (Envigo, Tampa, FL) for experiments involving HSV-GS11) were inoculated with the appropriate viral vector or phosphate-buffered saline (pH 7.3) (50 µl/mouse) on the lightly abraded plantar surfaces of both rear feet as previously described (40). To facilitate efficient uptake of vector, and to minimize the amount of abrasion required, the feet were saline-treated prior to infection. The following procedure was employed: mice were first anesthetized with isoflurane by inhalation. Flunixin meglumine (1.1 mg/kg) was then administered intramuscularly (IM) to alleviate any pain associated with the procedure. Thereafter, 25-50 µl (no more than 50µl) of sterile 10% saline were injected sub-epidermally under both rear footpads. The mice were then returned to their cages. Four h later, the mice were anesthetized by IM injection of 10-20 µl of a cocktail of acepromazine (2.5-3.75 mg/kg), xylazine (7.5-11.5 mg/kg) and ketamine (30-45 mg/kg). Both rear footpads of the anesthetized mice were then lightly abraded with an emery board to scratch the keratinized layer of the skin to allow the virus to adsorb efficiently. The anesthetized mice were rested on their backs, and 50 µl of the appropriate dilution of viral vector were placed on the footpads using a pipette. The viral vector was allowed to adsorb until the mice awoke. To activate vector replication, a combination of heat and mifepristone/ulipristal treatment was used. Heat treatment was performed by immersion of hindlimbs in a temperature-controlled water bath 3 h after virus administration. Mice were allowed to recover at 37°C for 15 min.

Mifepristone (0.5 mg/kg) or ulipristal (50 µg/kg) in DMSO was administered intraperitoneally (IP) at the time of immunization and, again, 24 h later.

### HSV-1 challenge in mice

Immunized and control mice were challenged by infection on both rear footpads with 10,000 pfu, typically, of HSV-1 strain 17*syn*+. The procedure for the saline pre-treatment, anesthesia and application of virus was the same as that described above for application of the viral vector, except that no mifepristone/ulipristal or heat treatment was administered. For the efficacy determination, a modified endpoint analysis was used. Mice were monitored daily (with cages coded in a masked fashion). When mice reached clinical endpoints indicating severe CNS infection (bilateral hindlimb paralysis, inability to move when touched, or trembling), they were euthanized.

### Blood collection, and serum and lymphocyte isolation

For serum neutralization studies, blood was collected by retro-orbital bleeding prior to immunization. For retro-orbital bleeding, mice were anesthetized with ketamine (90 mg/kg) and xylazine (10 mg/kg) IP. For the collection of serum and lymphocytes at 3 weeks after immunization, mice from each group were anesthetized by inhalation of 2-3% isoflurane. The total blood volume of each mouse was collected, and the mice were euthanized by cervical dislocation. PBMCs were isolated by Ficoll gradient separation using Lymphoprep (Miltenyi Biotec, Bergisch Gladbach. Germany) according to the manufacturer’s protocol.

### Neutralizing antibody assay

After collection, the blood was allowed to clot for 30 min. After centrifugation at 800 × g, the serum was collected. Serum samples were heated to 56°C for 1 h to inactivate complement and were then diluted 1:10 in complete DMEM containing 10% heat-inactivated FBS. For HSV-1 neutralization, 50 µl of a suspension containing approximately 100 pfu of HSV-1 strain 17*syn*+ were added to each dilution of serum to a final volume of 100 µl. For EIV neutralization, 50 µl of a suspension containing approximately 100 TCID50 units of EIV Prague/56 were added to each dilution of serum to a final volume of 100 µl. Initial serum dilution therefore was 1:20. Serum-virus mixtures were placed on a rocker at room temperature for 1 h, and the amount of virus that was not neutralized at a given concentration of serum was titrated on RS (HSV-1) or MDCK (EIV) cell monolayers in order to calculate the neutralizing antibody titers.

### Responder cell frequency (RCF) assay

HSV-1- or EIV-specific T cells were quantified by a modified limiting dilution lymphoproliferation assay (54), using either cell-free HSV-1 or EIV Prague/56 HA protein lysate and control antigen lysate. Briefly, wells of 96 well plates were coated with 20 µl/well of antigen extract (HSV-1 or RS cell control lysate; or EIV Prague/56 HA or Vero cell control lysate) and were allowed to air-dry in a laminar flow hood. Dilutions of mouse PBMCs in DMEM were added to each well such that each well contained a minimum of 1 and a maximum of 10 lymphocytes per well in a volume of 100 µl complete medium (with serum). The plates were then incubated at 37°C. After 24 h, medium containing 10 µCi of H3-thymidine was added to each plate for 12 hours, medium was replaced and the plates were incubated for an additional 72 h. The wells were harvested, and the DNA was precipitated in 20 volumes of cold 10% trichloroacetic acid, transferred onto glass fiber discs (Whatman GF/C) by filtration, rinsed with 95% ethanol, and dried using a heat lamp. The filters were then transferred to scintillation vials with Scintiverse (Fisher Scientific) and counted. The counts per minute (cpm) of H3-thymidine were converted to RCF using the maximum likelihood estimate method of Levin et al. (55).

### Viral DNA replication and transcription

Groups (n=5) of 4-6-week old female outbred Swiss Webster mice were anesthetized and inoculated with an HSV-GS vector on both rear footpads as described above. Ulipristal (50 µg/kg) was administered IP at the time of infection. Three h after virus administration, heat treatment was performed by immersion of rear feet in a 45°C water bath for 10 min. Mice were recovered at 37°C for 15 min. Mice were sacrificed 24 h after heat induction, and the feet were dissected and snap-frozen in RNAlater (Sigma-Aldrich). DNA and RNA were extracted by grinding the tissues in TRIzol (Thermo Fisher Scientific), and back-extracting the DNA from the interface. Extracted DNA was subjected to Taqman real-time PCR (qPCR) for quantitative analysis of HSV-1 DNA (using HSV-1 DNA polymerase primers/probe) (56). RNA was analyzed by reverse transcription and Taqman real-time PCR (RT-qPCR) for the presence of ICP4, thymidine kinase (TK), capsid protein VP19c and glycoprotein C (gC) transcripts. Normalization of DNA and RNA quantities was relative to the cellular gene APRT. Sequences of primers and probes used are provided in Table 3.

### EIV Prague/56 HA ELISA

In order to produce the antigen for the ELISA, the insert from plasmid pBS-EIVPrague56/HA (described above) was subcloned into expression vector pET-31b (Millipore), and the EIVPrague56/HA protein was expressed in *E. coli* in accordance with the manufacturer’s instructions. After induction and growth, the bacteria were lysed, proteins were isolated in the presence of protease inhibitors, and the proteins were used to coat a 96 well ELISA plate which was allowed to air-dry. Dilutions (1:10 to 1:200) of murine serum were applied to the wells, and the plate was incubated at room temperature for 30 minutes. The serum was removed, and the wells were washed 2 × with PBS. Horseradish peroxidase-conjugated anti-IgG antibody was added to the wells, the plate was incubated for 30 min at room temperature, and the wells were washed 2 × with PBS. 3,3’,5,5’-Tetramethylbenzidine was added to the wells, and the plate was incubated for 10 minutes and then read with a spectrophotometer plate reader. Results were evaluated based on the mean of sample to negative control mean (S/N) ratio.

### Statistical analyses

Data sets were analyzed using a two-way ANOVA analysis (no RM). Unless otherwise stated in the figure legends, data are presented as mean values with standard deviation. Kaplan-Meier survival curves were analyzed for significance using the log rank test. No data sets that were analyzed for statistical significance varied in sample size by more than 3-fold.

### Ethics statement

The animal studies were approved by the University of Florida Institutional Animal Care and Use Committee (IACUC) and were performed in accordance with the approved protocol (Protocol #0541). All experiments were carried out under adherence with the Guide for the Care and Use of Laboratory Animals of the National Institutes of Health and the American Veterinary Medical Association Guidelines on Euthanasia.

## ACKNOWLEDGMENTS

This work was supported by a contract from HSF Pharmaceuticals (to DCB) and an award from the UF Opportunity Fund (to DCB).

The authors also acknowledge technical assistance by Terri Edwards and Sean Taasan, and statistical analysis by Biaming Zho.

RV is the founder of HSF Pharmaceuticals SA, a company with an exclusive focus on research and early development activities.

